# The association between brain serotonin 2A receptor binding and neuroticism in healthy individuals: A Cimbi database independent replication study

**DOI:** 10.1101/2023.08.24.554575

**Authors:** Emma S. Høgsted, Vincent Beliveau, Brice Ozenne, Martin K Madsen, Claus Svarer, Vibeke NH. Dam, Annette Johansen, Patrick M. Fisher, Gitte Moos Knudsen, Vibe G Frokjaer, Anjali Sankar

**Author notes:** **Correspondence:** Anjali Sankar, PhD, Neurobiology Research Unit, Copenhagen University Hospital Rigshospitalet, 6-8 Inge Lehmanns Vej, Building 8057, 2100 Copenhagen O, Denmark. denotes shared authorship (Vibe G Frokjaer, Anjali Sankar).

## Abstract

**Background:** Using the [^18^F]altanserin tracer to image serotonin 2A receptors (5-HT_2A_R), we previously showed that there exists a positive association between cortical 5-HT_2A_R binding and the inward facets of neuroticism, namely depression, anxiety, self-consciousness, and vulnerability. Fairly recently, the [^11^C]Cimbi-36 tracer was also shown to be a suitable radioligand for imaging 5-HT_2A_ receptors in the human brain. In the present study, we examined whether our previously reported finding of the association between 5-HT_2A_R binding and the inward facets of neuroticism can be replicated in an independent sample of healthy individuals scanned using the newer [11C]Cimbi-36 tracer. Furthermore, to determine whether this association of 5-HT_2A_R binding with neuroticism merely reflects its known relation to stress-coping related indices such as cortisol dynamics. The present study also investigated the potential role of cortisol awakening response on the association between 5-HT_2A_R binding and the inward facets of neuroticism.

**Methods:** Sixty-nine healthy volunteers underwent a [^11^C]CIMBI-36 scan for the assessment of 5-HT_2A_R binding, completed the standardized NEO-PI-R personality questionnaire, and provided salivary samples for the determination of cortisol awakening response. A linear latent variable model (LVM) was used to examine the association between 5-HT_2A_R binding and the inward facets of neuroticism with adjustment for age, sex, cortisol awakening response, and MR scanner. A second latent variable model examined the potential moderating effect of cortisol awakening response on the association between 5-HT_2A_R binding and the inward facets of neuroticism.

**Results:** We replicated a positive association between 5-HT_2A_R binding and the inward facets of neuroticism (r=0.37, p=0.015). We saw no moderating effect of the cortisol awakening response on this association (p=0.98).

**Conclusions:** In an independent cohort of healthy individuals imaged with the [^11^C]CIMBI-36 tracer, we confirm the link between serotonin 2A receptor binding and the inward-directed facets of neuroticism that is independent of cortisol dynamics.

## INTRODUCTION

Several lines of evidence, including molecular imaging, psychopharmacology, and peripheral blood markers, strongly support the role of serotonin (5-HT) in the pathophysiology of depression.^1, 2^ Of the different 5-HT receptor systems, the serotonin 2A receptor (5HT_2A_R) system has been shown to be important for regulating functions that are disrupted in depression such as mood and cognition.^3^ In addition, 5-HT_2A_R is associated with certain personality traits that are robust risk factors for developing depression. For instance, our group was the first to show a positive association between 5-HT_2A_R binding and neuroticism in healthy individuals using positron emission tomography (PET) imaging with the [^18^F]altanserin tracer.^4^ Furthermore, we showed that this link between serotonergic function and neuroticism was stronger in individuals at high familial risk for developing depression relative to individuals at low familial risk.^5^ Interestingly, in both datasets the facets or dimensions of neuroticism that were closely linked to 5-HT_2A_R binding were those facets that are directed “inward” such as depression, anxiety, self-consciousness, and vulnerability to stress, as opposed to the “outward” directed facets such as impulsivity and angry hostility.^5^

Along with [^18^F]altanserin, [^11^C]Cimbi-36 has also been shown to be a suitable radioligand for imaging 5-HT_2A_receptors in the human brain, albeit more recently.^6^ Although both altanserin and Cimbi-36 have high affinity for the 5-HT_2A_R, the two compounds have distinct pharmacological properties. For instance, altanserin is an antagonist^7^ while Cimbi-36 is an agonist at the 5-HT_2A_R.^8^ Cimbi-36 has similar affinity in vitro for binding to 5-HT_2A_ and 5-HT_2C_receptors^8^ whereas altanserin is more selective for binding to 5-HT_2A_than to 5-HT_2c_ receptors.^9^ Additionally, the PET radioligands [^18^F]altanserin and [^11^C]Cimbi-36 are radiolabeled with different radioisotopes. Moreover, [^11^C]Cimbi-36 PET scanning is performed with a bolus injection,^6^ while a steady-state approach with bolus followed by constant infusion is required for the [18F]altanserin scan.^10^

As the previous study of the association between 5-HT_2A_R and the inward facets of neuroticism was performed using [^18^F]altanserin, we aimed to investigate whether this association could be replicated with [^11^C]Cimbi-36 PET imaging in an independent group of patients. Methodological inconsistencies, importantly differences in tracers are often cited as reasons for mixed findings in the PET literature. Thus, a study of whether the finding can be replicated in an independent sample of healthy individuals using the newer [^11^C]Cimbi-36 radiotracer is warranted. Furthermore, it is unclear whether the association between and neuroticism is a mere reflection of the known association between variations in serotonergic measures and stress-coping indices such as cortisol dynamics.^11, 12^ In our previous studies, we could not examine the potential role of cortisol dynamics as we did not have these data.^4^

Therefore, in the present study, we include cortisol markers to examine if the known associations between the 5-HT_2A_R binding and the inward facets of neuroticism are moderated by stimulated cortisol response in terms of the cortisol awakening response.

## METHODS

### Participants

We sought to identify individuals within the CIMBI database that met the following criteria: (a) completed a baseline [^11^C]CIMBI-36 scan, (b) completed an assessment of neuroticism using the NEO-PI-R (c) provided salivary samples for the determination of cortisol levels, and (d) were without a lifetime history of neurological and psychiatric disorders. Sixty-nine healthy individuals met this inclusion criteria and were included in the present study. The individuals included herein participated in research studies at the Neurobiology Research Unit, Copenhagen University Hospital (Rigshospitalet), Copenhagen between 2012-2021.

Written informed consent was obtained from all participants before their inclusion in the studies conducted at the Neurobiology Research Unit. The individual study protocols were approved by the Ethics Committee of Copenhagen and Frederiksberg, Denmark.

### Procedure

#### MR Imaging

The MR-scans were performed on a 3T Verio scanner (n=26) and a 3T Prisma scanner (n=43) using the following parameters: verio/prisma: repetition time= 1900 ms, echo time= 2.32/2.58 ms, inversion time= 900 ms, flip angle= 9°, in-plane matrix= 256 × 256, in-plane resolution= 0.9 × 0.9 mm, 224 slices and a slice thickness of 0.9 mm, no gap. The images were segmented using VBM8 in SPM8^13^ to mask gray matter in the subsequent extraction of tissue time-activity curves.

#### PET data acquisition

Procedures for PET acquisition have been described in detail elsewhere.^14^ Briefly, scans were conducted using a high-resolution research tomography HRRT PET scanner with parameters: matrix=256 × 256 × 207 voxels, voxel size=1.22 × 1.22 × 1.22 mm. All participants underwent a ten-minute transmission scan and underwent a 120-minute emission scan which started at the time of the intravenous bolus injection of [^11^C]Cimbi-36 (mean injected activity: 499 ± 109 MBq; injected cold Cimbi-36 dose: 0.72 ± 0.44 µg). [^11^C]Cimbi-36 scanning data were reconstructed into 45 dynamic frames (6 × 10 s, 6 × 20 s,6 × 60 s, 8 × 120 s, and 19 × 300 s).

#### PET processing and quantification

Motion correction of the PET images was performed using AIR (version 5.2.5). All frames were aligned to the first 5-minute frame, and partial-volume correction was not applied. For each subject the 3D T1-weighted MPRAGE image was co-registered to the PET image using SPM8. Delineation of regions of interest was performed using PVE-lab,^15^, and performed on the individual’s T1-weighted MPRAGE image. The mean tissue time activity curves for the set of grey matter voxels within a region of interest (ROI) were extracted for kinetic modeling. A global neocortical region was defined as a volume-weighted mean of the following cortical ROIs: orbitofrontal cortex, superior, middle, and inferior frontal gyrus; superior, middle, and inferior temporal gyrus, sensorimotor cortex, parietal cortex, and occipital cortex. A composite neocortex ROI was used because the signal is very highly correlated between the neocortical regions.^16^

Kinetic modeling using the simplified reference tissue model (SRTM) ^17^ with the cerebellum (excluding vermis) was performed with PMOD software version 3.0 (PMOD, Zurich, Switzerland). The calculated non-displaceable binding potential (BP_ND_) served as the outcome measure for quantifying 5-HT_2A_ binding. Mean BP_ND_ from the neocortex was extracted. BP_ND_ is defined as:

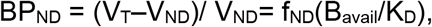

where V_T_ and V_ND_ are the distribution volumes in the ROI and reference region (cerebellum), respectively, from two-tissue compartment modeling with arterial input measurements as previously described.^14^ Cerebellum has been previously validated as an appropriate reference region for [^11^C]Cimbi-36.^14^

#### Neuroticism

Participants completed the Danish version of the 240-item NEO Personality Inventory-Revised (NEO-PI-R).^18^ NEO-PI-R, a self-reported personality questionnaire, evaluates the broad personality dimensions of neuroticism, extraversion, openness, agreeableness, and conscientiousness. Each dimension score is derived by summing the scores from the assessment of six constituent personality traits or facets namely, anxiety, depression, self-conscientiousness, vulnerability, angry hostility, and impulsiveness, and each trait score is derived by summing the scores from eight constituent items. There are no meaningful cut-offs for the NEO-PI-R scale as it captures variations in personality traits in the healthy spectrum.

The outcome measure of neuroticism for this study were the constituent traits of anxiety, depression, self-conscientiousness, and vulnerability to stress as these were facets of neuroticism that were found to be associated with 5-HT_2A_R binding in an earlier study of individuals with a familial risk of developing depression.^19^

#### Cortisol Awakening Response

The cortisol awakening response (CAR), i.e., the dynamic increase in cortisol levels that occurs within the first hour upon morning awakening was determined as a marker stimulated stress hormone response. For the assessment of CAR, participants were instructed to collect saliva samples immediately after awakening and after 15, 30, 45, and 60 minutes. Saliva samples were collected in Salivette tubes (Sarstedt, Neubringen, Germany). Participants were told to avoid food, drinking, brushing their teeth, and smoking during the first hour after waking up. If saliva samples were collected more than 10 minutes after awakening, the dataset was not included in our analyses and was categorized as non-compliant. The saliva samples collected at home were stored in the refrigerator and returned to the laboratory the day after completion of sampling and if collected during the weekend, after a maximum of three days after collection. The CAR measurements were computed as the area under the curve with respect to increase from baseline at awakening (AUCi) indicating reactivity of the Hypothalamic Pituitary Adrenal (HPA) axis. The saliva samples were analyzed in four different batches across the collection period to minimize experimental noise. Participant training, instructions, home-sampling procedures, storing and cortisol analyses were carried out as described in Frokjaer et al.^20^

### Statistical analyses

#### *Association between neocortical* 5-HT_2A_R *binding and neuroticism*

We applied a linear latent variable model (LVM) using the *lava* package^21^ implemented in R to first examine the association between 5-HT_2A_R binding in the neocortex and neuroticism in healthy controls. LVMs are flexible structural equation models, which permit the modeling of complex hierarchical structures and summarizing multivariate data into a single latent variable, enabling ease of interpretation as well as reducing issues related to multiple testing of correlated variables. The PET variable was log-transformed regional BP_ND_ values from the neocortex, and the neuroticism variables were scores on anxiety, depression, self-conscientiousness, and vulnerability. The four constituent traits represent inward-directed facets of neuroticism. One latent variable representing the inward facets of neuroticism was constructed (Inward-Neuroticism_LV_). Age, sex, and AUCi were included as covariates on both Inward-Neuroticism_LV_ and log-transformed regional BP_ND_ values. MR Scanner type was included as an additional covariate on the log-transformed regional BP_ND_ values.

Associations are described both in terms of covariance coefficients (“*parameter estimate”*) and Pearson’s correlation coefficients (*r*). The moderating effect of AUCi on the association between 5-HT_2A_R binding and neuroticism was evaluated in a separate LVM. For this analysis, AUCi values were split into three equal groups with low, medium, and high values of AUCi. A third LVM examined the moderating effect of sex on the association between 5HT2A binding and neuroticism. Additional model paths were considered for all three LVMs using the *modelsearch* function within *lava* and additional paths were added if the statistical significance of the Rao score test for the individual path was *p*_*fwe*_<.05, adjusting for all possible paths. This procedure was repeated until no new model paths were supported.

## RESULTS

### Demographic, Clinical, Scanner, and Radiotracer Information

Demographics and radiotracer information for healthy individuals are detailed in Table 1. There was a slight over-representation of men in this sample (58%), but there was no significant difference in age (*p*=0.92), BMI (*p*=0.63), or weight-adjusted [^11^C]Cimbi-36 injected mass (*p*= 0.11) between men and women.

**Table 1:** Demographic, Clinical, Scanner and Radiotracer Information.

| Healthy Individuals (n=69) |  |  |
| --- | --- | --- |
|  | Mean | SD |
| Age (years) | 28.6 | 7.8 |
| BMI (kg/m<sup>2</sup>) | 23.9 | 3.0 |
| MDI | 4.7 | 3.2 |
| AUCi | 66.6 | 272.3 |
| Minutes from awakening to CAR | 1.8 | 3.4 |
| Injected tracer dose (MBq) | 0.01 | 0.006 |
|  | N | % |
| Women | 29 | 42 |
| MRI scanner (3T Verio) | 26 | 38 |

Abbreviations: SD: standard deviation; MDI: major depressive inventory; AUCi: area under the curve with respect to increase from baseline at awakening (AUCi); CAR: cortisol awakening response.

### Model Summary

All four constituent sub-facets of neuroticism loaded well onto the latent variable Inward-Neuroticism_LV_ (estimate range = 0.81:1.42, *p*< 0.001). Score tests supported an additional partial correlation between the depression sub-facet of neuroticism and sex (*p*_*fwe*_=0.04). The first LVM-based model showed a positive association between 5-HT_2A_ receptor BP_ND_ and the inward facets of neuroticism (summarized by the inward-neuroticism_LV_) across all participants (parameter estimate = 0.13, 95% CI=0.025: 0.23, *r*=0.37, *p*=0.015, Figures 2). Estimation of effects of covariates revealed a significant association between AUCi and 5-HT_2A_ receptor BP_ND_ (parameter estimate = 8.51, 95% CI=0.41: 16.62, r=8.51, *p*=0.04). Additionally, there were significant negative associations of age with both 5-HT_2A_ receptor BP_ND_(parameter estimate = -0.011, 95% CI=-0.015: -0.007, *p*<0.001), and inward-neuroticism_LV_ (parameter estimate = -0.10, 95% CI=-0.2: -0.001, *p*=0.048). Women had greater 5-HT_2A_ receptor BP_ND_ than men (parameter estimate = 0.10, 95% CI=0.04: 0.16, *p*=0.001).

**Figure 1:**
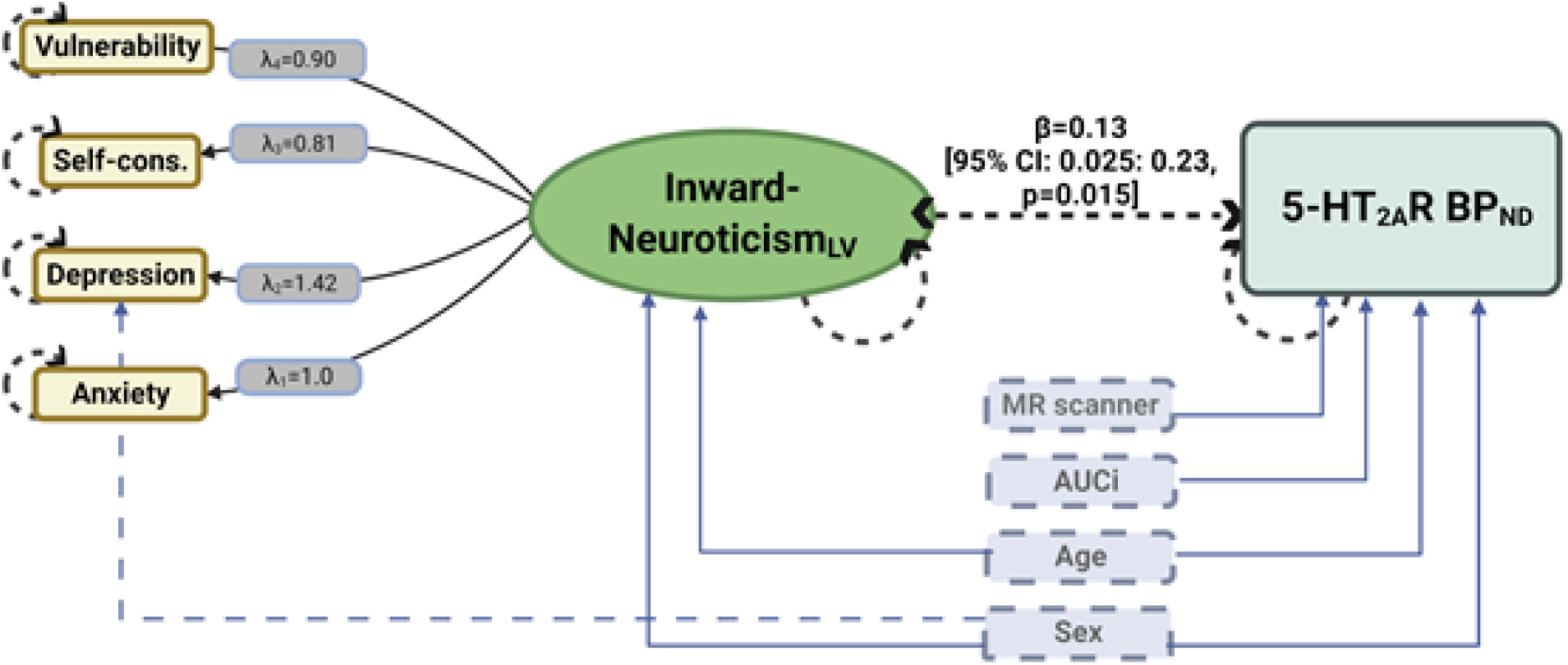
Latent variable model to estimate the association between 5-HT2AR binding and the inward facets of neuroticism. Figure 1 shows the estimated latent variable model for the association between 5-HT_2A_R binding and the inward facets of neuroticism. The green ellipse represents the latent variable inward-neuroticism_LV_. The dashed line between depression and sex represents an additional shared correlation. Circular dashed lines represent error estimates included in the model. *λ* represents the loading of a given neuroticism sub-facet estimate onto its respective latent variable. Both inward-neuroticism_LV_ and neocortex BP_ND_ were adjusted for age and sex, and neocortex BP_ND_ was additionally adjusted for AUCi (cortisol awakening response measured as area under the curve concerning increase from baseline at awakening) and MR scanner type.

**Figure 2:**
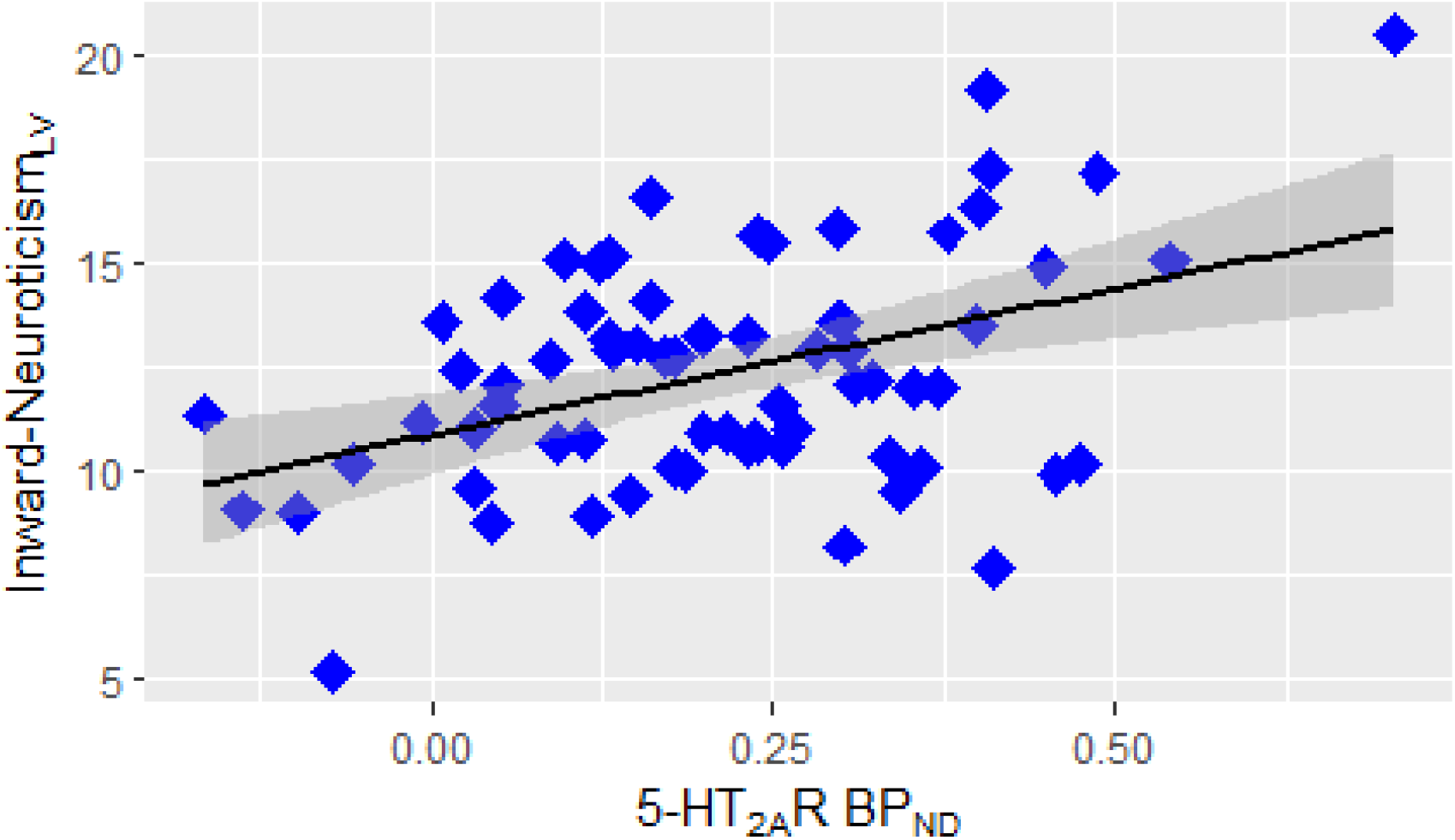
Positive association between 5-HT_2A_R binding in the neocortex and the inward facets of neuroticism in healthy controls. Figure2 shows positive association between latent estimates of inward-neuroticism (summarized as Inward-Neuroticism_LV_) 5-HT_2A_R binding in the neocortex across (log transformed BP_ND_ scores) all participants (r=0.37, p=0.015) obtained from the latent linear latent variable model. Both inward-neuroticism_LV_ and neocortex BP_ND_ were adjusted for age and sex, and neocortex BP_ND_ was additionally adjusted for AUCi (cortisol awakening response measured as area under the curve concerning increase from baseline at awakening) and MR scanner type. The black line represents the overall slope and the grey shading represents 95% confidence intervals.

There was no moderating effect of either AUCi (p=0.98) or sex (p=0.60) on the association between 5-HT_2A_ receptor BP_ND_ and inward-neuroticism_LV_.

## DISCUSSION

This study is an independent replication of our previously reported finding of a positive association between 5-HT_2A_R binding and the inward facets of neuroticism. Using latent variable models, we showed a positive association between 5-HT_2A_ receptor BP_ND_ and the inward facets of neuroticism across all participants. Furthermore, there was no evidence that either sex or cortisol levels moderated this association.’

Using a different tracer to image 5-HT_2A_ receptors, namely [^18^F]altanserin, we previously showed a significant association between 5-HT_2A_R binding and the inward facets of neuroticism in healthy individuals.^4^ Subsequently, using the same tracer, we found that this association was stronger in individuals with high-familial risk for depression relative to those with low-familial risk.^5^ The current replication of the association between serotonergic function and the inward facets of neuroticism in an independent sample with the [^11^C]Cimbi-36 tracer provides further support to the notion that serotonergic neurotransmission contributes to the individual differences in personality risk factors of depression.

Several lines of investigation have pointed to altered 5-HT_2A_R levels in major depression. For instance, neuroimaging studies have largely shown high 5-HT_2A_ receptor levels in patients with depression relative to healthy individuals.^22, 23^ 5-HT_2A_R stimulates cortisol secretion,^24^ another marker of vulnerability to depression.^25^ In the present study, 5-HT_2A_R was significantly associated with cortisol awakening response (as measured by AUCi) in addition to the inward facets of neuroticism. These findings are also corroborated by genetic studies which show that a common polymorphism of the 5-HT_2A_R gene (1438A/G) is associated with higher cortisol awakening response and higher neuroticism. ^24^Together, these findings suggest that the 5-HT_2A_R may be a stress response system.^26, 27^

We did not find evidence for the association between cortisol levels (AUCi) and neuroticism levels. Although their association has been previously reported, findings tend to be inconsistent,^28, 29^ and reported mostly in women.^29^ Additionally, this study also found a negative association between 5-HT_2A_R binding and age, a finding previously also observed with altanserin.^30^ Sex differences in 5-HT_2A_R binding have not been previously reported, and therefore, need further study.

The findings of the study should be considered in the context of its limitations. 5-HT_2A_R is most abundantly found in the neocortex where the signal with the applied PET method is also best captured, and therefore we limited our investigation to neocortical 5-HT_2A_R binding and its association with the inward facets of neuroticism. Whether there is a region-specific difference in the association is unclear. We did not include individuals older than 60 years, therefore our results may not be generalizable to older populations

In conclusion, in this dataset comprising molecular imaging with [^11^C]Cimbi-36 and personality measures, we confirm the link between 5-HT_2A_R binding and the inward-directed personality facets of neuroticism. This association does not appear to be dependent on cortisol dynamics, and persists across sex. Whether 5-HT_2A_R binding alterations lead to high neuroticism, or whether shared genetic factors predispose individuals to altered 5-HT_2A_R binding and high neuroticism remains to be studied.

## REFERENCES

1. Jauhar S, Cowen PJ, Browning M. Fifty years on: Serotonin and depression. Journal of Psychopharmacology 2023; 37(3): 237–241.

2. Jauhar S, Arnone D, Baldwin DS, Bloomfield M, Browning M, Cleare AJ et al. A leaky umbrella has little value: evidence clearly indicates the serotonin system is implicated in depression. Molecular Psychiatry 2023: 1–4.

3. Landolt HP, Wehrle R. Antagonism of serotonergic 5-HT2A/2C receptors: mutual improvement of sleep, cognition and mood? European Journal of Neuroscience 2009; 29(9): 1795–1809.

4. Frokjaer VG, Mortensen EL, Nielsen FÅ, Haugbol S, Pinborg LH, Adams KH et al. Frontolimbic serotonin 2A receptor binding in healthy subjects is associated with personality risk factors for affective disorder. Biological psychiatry 2008; 63(6): 569–576.

5. Frokjaer VG, Vinberg M, Erritzoe D, Baaré W, Holst KK, Mortensen EL et al. Familial risk for mood disorder and the personality risk factor, neuroticism, interact in their association with frontolimbic serotonin 2A receptor binding. Neuropsychopharmacology 2010; 35(5): 1129–1137.

6. Ettrup A, da Cunha-Bang S, McMahon B, Lehel S, Dyssegaard A, Skibsted AW et al. Serotonin 2A receptor agonist binding in the human brain with [11C] Cimbi-36. Journal of Cerebral Blood Flow & Metabolism 2014; 34(7): 1188–1196.

7. Kristiansen H, Elfving B, Plenge P, Pinborg LH, Gillings N, Knudsen GM. Binding characteristics of the 5-HT2A receptor antagonists altanserin and MDL 100907. Synapse 2005; 58(4): 249–257.

8. Ettrup A, Hansen M, Santini MA, Paine J, Gillings N, Palner M et al. Radiosynthesis and in vivo evaluation of a series of substituted 11 C-phenethylamines as 5-HT 2A agonist PET tracers. European journal of nuclear medicine and molecular imaging 2011; 38: 681–693.

9. Herth MM, Kramer V, Piel M, Palner M, Riss PJ, Knudsen GM et al. Synthesis and in vitro affinities of various MDL 100907 derivatives as potential 18F-radioligands for 5-HT2A receptor imaging with PET. Bioorganic & medicinal chemistry 2009; 17(8): 2989–3002.

10. Pinborg LH, Adams KH, Svarer C, Holm S, Hasselbalch SG, Haugbøl S et al. Quantification of 5-HT2A receptors in the human brain using [18F] altanserin-PET and the bolus/infusion approach. Journal of Cerebral Blood Flow & Metabolism 2003; 23(8): 985–996.

11. Way BM, Taylor SE. The serotonin transporter promoter polymorphism is associated with cortisol response to psychosocial stress. Biological psychiatry 2010; 67(5): 487–492.

12. O’hara R, Schröder C, Mahadevan R, Schatzberg A, Lindley S, Fox S et al. Serotonin transporter polymorphism, memory and hippocampal volume in the elderly: association and interaction with cortisol. Molecular psychiatry 2007; 12(6): 544–555.

13. Ashburner J, Barnes G, Chen C, Daunizeau J, Flandin G, Friston K et al. SPM8 manual. Functional Imaging Laboratory, Institute of Neurology 2012.

14. Ettrup A, Svarer C, McMahon B, da Cunha-Bang S, Lehel S, Møller K et al. Serotonin 2A receptor agonist binding in the human brain with [11C] Cimbi-36: test–retest reproducibility and head-to-head comparison with the antagonist [18F] altanserin. Neuroimage 2016; 130: 167–174.

15. Svarer C, Madsen K, Hasselbalch SG, Pinborg LH, Haugbøl S, Frøkjær VG et al. MR-based automatic delineation of volumes of interest in human brain PET images using probability maps. Neuroimage 2005; 24(4): 969–979.

16. Spies M, Nasser A, Ozenne B, Jensen PS, Knudsen GM, Fisher PM. Common HTR2A variants and 5-HTTLPR are not associated with human in vivo serotonin 2A receptor levels. Human Brain Mapping 2020; 41(16): 4518–4528.

17. Lammertsma AA, Hume SP. Simplified reference tissue model for PET receptor studies. Neuroimage 1996; 4(3 Pt 1): 153–158.

18. Skovdahl-Hansen H, Hk MES. Dokumentation for den danske udgave af NEO-PI-R og NEO-PI-R Kort Version. 2004. Copenhagen, Denmark, Dansk Psykologisk Forlag Ref Type: Report.

19. Frokjaer VG, Vinberg M, Erritzoe D, Baaré W, Holst KK, Mortensen EL et al. Familial risk for mood disorder and the personality risk factor, neuroticism, interact in their association with frontolimbic serotonin 2A receptor binding. Neuropsychopharmacology 2010; 35(5): 1129–1137.

20. Frokjaer VG, Erritzoe D, Holst KK, Jensen PS, Rasmussen PM, Fisher PM et al. Prefrontal serotonin transporter availability is positively associated with the cortisol awakening response. European Neuropsychopharmacology 2013; 23(4): 285–294.

21. Holst KK, Budtz-Jørgensen E. Linear latent variable models: the lava-package. Computational Statistics 2013; 28(4): 1385–1452.

22. Hrdina PD, Demeter E, Vu TB, Sótónyi P, Palkovits M. 5-HT uptake sites and 5-HT2 receptors in brain of antidepressant-free suicide victims/depressives: increase in 5-HT2 sites in cortex and amygdala. Brain research 1993; 614(1-2): 37–44.

23. McKeith I, Marshall E, Ferrier I, Armstrong M, Kennedy W, Perry R et al. 5-HT receptor binding in post-mortem brain from patients with affective disorder. Journal of affective disorders 1987; 13(1): 67–74.

24. Fiocco AJ, Joober R, Poirier J, Lupien S. Polymorphism of the 5-HT2A receptor gene: Association with stress-related indices in healthy middle-aged adults. Frontiers in Behavioral Neuroscience 2007; 1: 83.

25. Dedovic K, Ngiam J. The cortisol awakening response and major depression: examining the evidence. Neuropsychiatric disease and treatment 2015: 1181–1189.

26. Murnane KS. Serotonin 2A receptors are a stress response system: implications for post-traumatic stress disorder. Behavioural pharmacology 2019; 30(2-): 151.

27. Leonard BE. The HPA and immune axes in stress: the involvement of the serotonergic system. European Psychiatry 2005; 20(S3): S302–S306.

28. Sundin ZW, Chopik WJ, Welker KM, Ascigil E, Brandes CM, Chin K et al. Estimating the associations between big five personality traits, testosterone, and cortisol. Adaptive Human Behavior and Physiology 2021: 1–34.

29. Oswald LM, Zandi P, Nestadt G, Potash JB, Kalaydjian AE, Wand GS. Relationship between cortisol responses to stress and personality. Neuropsychopharmacology 2006; 31(7): 1583–1591.

30. Adams KH, Pinborg LH, Svarer C, Hasselbalch SG, Holm S, Haugbøl S et al. A database of [18F]-altanserin binding to 5-HT2A receptors in normal volunteers: normative data and relationship to physiological and demographic variables. Neuroimage 2004; 21(3): 1105–1113.

